# Puromycin is incorporated into regenerating flagella of *Chlamydomonas reinhardtii* as an indicator of nascent flagellar proteins

**DOI:** 10.1101/2022.01.27.478094

**Authors:** Tomohiro Kubo, Natsumi Kanou, Toshiyuki Oda

## Abstract

The unicellular alga *Chlamydomonas reinhardtii* assembles flagella after pH shock-induced deflagellation, in which process the cell vigorously synthesizes flagellar proteins. Previously, newly synthesized flagellar proteins were chased using radioisotopes. Here, we have developed an alternative, non-radioactive method using the surface sensing of translation (SUnSET) assay, an assay that takes advantage of the incorporation of the antibiotic puromycin into newly synthesized peptides. Just after deflagellation, puromycin-labeled proteins increased dramatically in the cytoplasm and newly assembled flagella. Axonemal incorporation of newly synthesized proteins occurred from the distal tip of the flagellum. Cycloheximide, a protein synthesis inhibitor, almost completely prevented the cellular and flagellar incorporation of puromycin, although, as reported previously, it allowed reassembly of half-length flagella from the cytoplasmic “flagellar precursor” proteins. In contrast to wild-type cells, a cycloheximide-resistant mutant *act2* produced nearly full-length flagella containing puromycin-labeled proteins even in the presence of cycloheximide. These and other results demonstrate that the SUnSET method serves as a powerful tool for visualizing flagellar incorporation of nascent proteins in *Chlamydomonas.*

## Introduction

Building cellular organelles, such as cilia and flagella (interchangeable terms), requires spatiotemporal regulation of protein synthesis and transport. The flagellum, a highly conserved whip-like cell organelle, is involved in various functions including cell motility, fluid flow generation, and signal transduction (Huangfu et al, 2003; Corbit et al, 2005). To build a well-ordered structure of flagellum requires a dynamic system called the intraflagellar transport (IFT), a bidirectional movement of the IFT particles along the axonemal microtubules driven by the molecular motors (Kozminski et al, 1993; Rosenbaum and Witman, 2002).

*Chlamydomonas reinhardtii,* a photosynthetic unicellular organism with two flagella per cell, serves as an ideal tool to study flagellar assembly. The cells detach them upon mechanical or pH shock but can reassemble new flagella to the original length of ~12 μm within two to three hours (Rosenbaum et al, 1969; Lefebvre et al, 1978). However, in the presence cycloheximide, a protein synthesis inhibitor, the cells can only produce flagella of ~6 μm (Rosenbaum et al, 1969). This observation has been interpreted as an indication that the steady-state cell possesses “flagellar precursor” sufficient to build half-length flagella and that the remaining half is completed using proteins synthesized after deflagellation.

The profile of protein synthesis after deflagellation has generally been studied using radioactive amino acid (Rosenbaum and Child, 1967; Rosenbaum et al, 1969; Lefebvre et al, 1978; Song and Dentler, 2001). About a decade ago, a nonradioactive method termed the <u>su</u>rface <u>se</u>nsing of translation (SUnSET) was reported (Schmidt et al, 2009). SUnSET employs an aminonucleoside antibiotic puromycin produced by *Streptomyces alboniger.* As a structural analog of the 3’ end of aminoacyl-tRNA, puromycin molecules are incorporated into nascent polypeptides and stop translation. The incorporated puromycin enables monitoring of the newly synthesized polypeptides by a monoclonal antibody against puromycin (Schmidt et al, 2009).

Despite its application for detecting protein synthesis in a broad range of situations, the utility of SUnSET during flagellar assembly was not reported before. Here, we applied SUnSET to the *Chlamydomonas* cells during flagellar regeneration and found that the puromycin-labeled proteins are detected in both cytoplasm and flagella. Therefore, the SUnSET method can be applied to this alga to investigate flagellar assembly and maintenance.

## Results and discussion

### Flagellar amputation induces de novo synthesis of flagellar proteins as assessed by the SUnSET method

During flagellar assembly, *Chlamydomonas* cells synthesize new flagellar proteins (Rosenbaum et al, 1969). We tested if the <u>su</u>rface <u>se</u>nsing of translation (SUnSET) method can be applied to chase newly synthesized proteins. SUnSET employs the antibiotic puromycin, a structural analog of the 3’ end of aminoacyl-tRNA, to label nascent polypeptides. Because puromycin is generally used to inhibit protein synthesis (Nathans, 1964), we first asked if the addition of puromycin affects cellular function. Up to 50 μg/ml puromycin did not cause any observable defects in flagellar growth (Figure 1A) and motility (data not shown). However, puromycin slightly retarded flagellar regeneration at 0.25 mg/ml, and caused production of half-length flagella at 1 mg/ml. This result is qualitatively similar to that of the addition of another protein synthesis inhibitor, cycloheximide (Figure 1A). Puromycin thus does not affect flagellar regeneration and function at concentrations <50 μg/ml, although it most likely inhibits protein synthesis at high concentrations.

**Figure 1.**
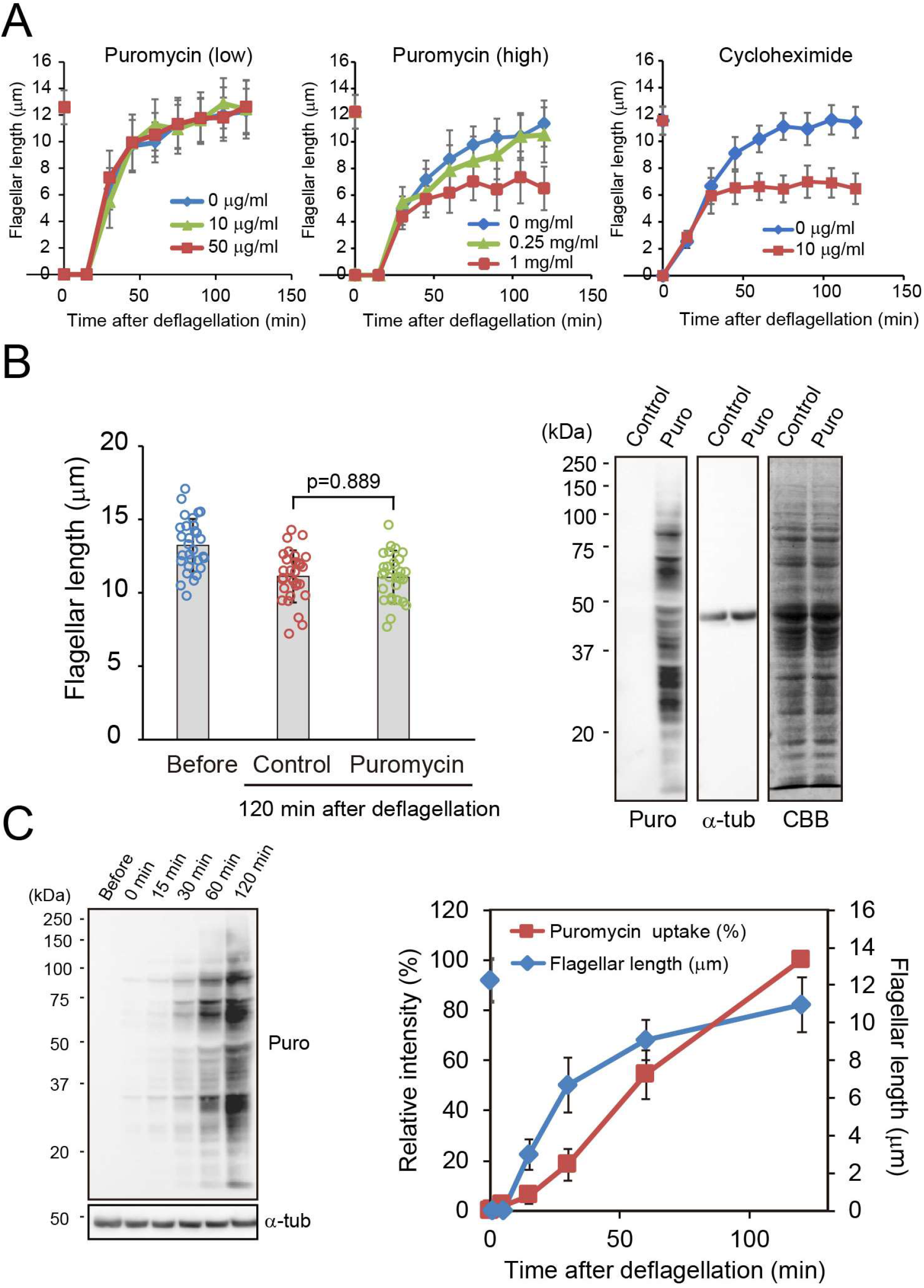
Puromycin labeling of proteins newly synthesized after deflagellation. (A) Flagellar regeneration in wild type in the presence of puromycin (left and middle) or cycloheximide (right). The pH of the medium was set to 4.5 to induce deflagellation and neutralized at time 0. The flagellar lengths before deflagellation are plotted on the ordinate. (B) Cellular incorporation of puromycin. Left panel: flagellar lengths before and 120 min after acid-induced deflagellation with or without puromycin. Right panel: Western blotting and CBB stained gels of whole cell lysates. The samples were probed with anti-puromycin (12D10) and α-tubulin antibodies. (C) Left panel: cellular accumulation of puromycin during flagellar regeneration. Right panel: flagellar regeneration kinetics and puromycin uptake of the cells.

To examine whether puromycin can be incorporated into the cell when new proteins are being synthesized, cells undergoing flagellar regeneration were treated with puromycin and processed for Western blotting. As expected, while cells without puromycin treatment showed no signal, puromycin-treated cells exhibited multiple clear bands, indicating that puromycin was incorporated into newly synthesized polypeptides (Figure 1B). Consistent with the previous finding that nascent flagellar proteins progressively accumulate in the cell during flagellar assembly (Lefebvre et al, 1978), the signals of puromycin-labeled proteins increased with time after deflagellation (Figure 1C).

### Puromycin-labeled proteins are incorporated into regenerating flagella

We then assessed whether the puromycin-labeled proteins can be incorporated into the flagellum. Nuclear-flagellar apparatuses (NFAps) were isolated from the cells treated or not treated with puromycin during flagellar regeneration and observed by immunofluorescence microscopy. The NFAps from puromycin-treated cells displayed prominent signals along flagella, whereas the NFAps from the cells without puromycin treatment showed essentially no signal (Figure 2A). Although the cellular localization of puromycin-labeled proteins was somewhat variable, we frequently observed strong signals in the basal body region. This observation is consistent with the idea that the newly synthesized proteins are initially accumulated at the basal body and thereafter transported into the flagellum.

**Figure 2.**
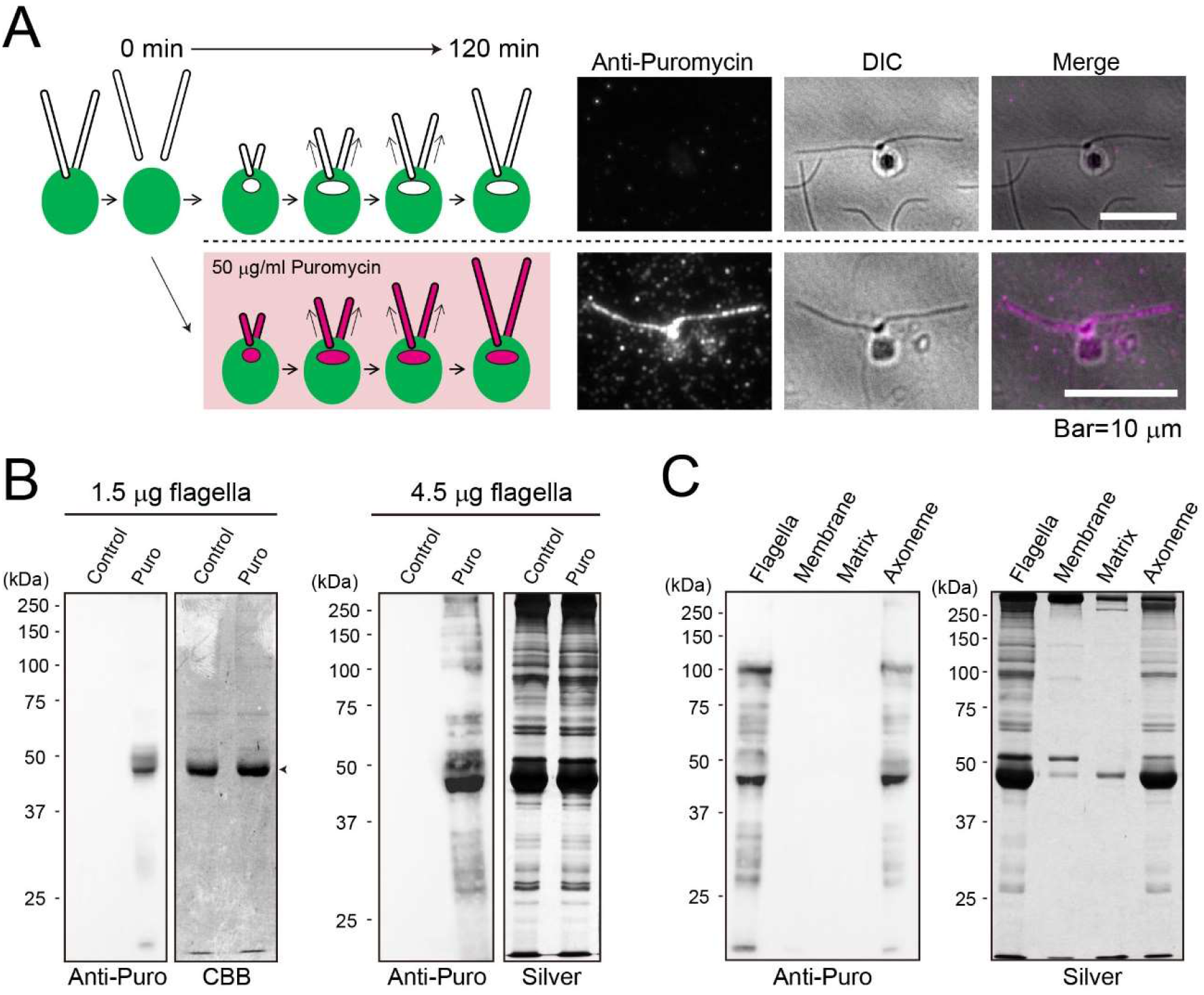
Puromycin-labeled proteins incorporated into regenerating flagella. (A) Indirect immunofluorescence microscopy of wild-type cells using anti-puromycin antibody. Left panels: schematic illustration of the time course. Right panels: fluorescence and DIC images at 120 min. Deflagellated cells were allowed to regenerate flagella in the absence (upper panels) or presence (lower panels) of puromycin for 2 hours. (B) Western blotting and CBB-or silver-stained gels of flagella isolated from cells treated with or without puromycin. (C) Western blotting and sliver-stained gels for different flagellar compartments.

As it is likely that puromycin-labeled proteins are incorporated into the flagellum, we biochemically examined isolated flagella. Regenerated flagella were isolated from cells that had been either treated or not treated with puromycin, and subjected to Western blotting using anti-puromycin antibody. As predicted, flagellar proteins from untreated cells showed no obvious bands (Figure 2B, left); in contrast, flagellar proteins from puromycin-treated cells gave rise to a prominent band, which most likely corresponded to α- and β-tubulins (~50 kDa). Western blotting using larger amounts of flagellar proteins displayed multiple bands, which mostly corresponded to the bands in silver-stained gels (Figure 2B, right). This suggests that a large fraction of flagellar proteins are labeled by puromycin albeit we do not yet know how much fraction underwent labeling. Western blotting of the membrane, matrix, and axonemal proteins from the same amounts of flagella revealed that the axonemal fraction accounts for most of the puromycin-labeled proteins (Figure 2C).

### Newly synthesized proteins are incorporated from the tip of the flagellum

In a growing flagellum, most structural proteins are incorporated at the distal tip (Johnson and Rosenbaum, 1992; Piperno et al, 1996; Marshall and Rosenbaum, 2001). To observe the location along the flagellum where nascent proteins are first incorporated, we localized the puromycin-labeled proteins during flagellar assembly. Different timings of puromycin treatment were tested (Figure 3, left). Cells treated with puromycin for 120 min immediately after the pH shock showed clear staining of the entire flagella as predicted (Figure 3, first row). The same was true for cells treated with puromycin for 60 min after the pH shock and then cultured in puromycin-free medium (Figure 3, 4^th^ row). However, cells treated with puromycin for 60 min from 60 min after the pH shock exhibited signals only at the distal tips (Figure 3, 2^nd^ row). Conversely, cells treated with puromycin only for 30 min immediately after the pH shock showed intensive staining in the proximal region but only faint staining in the distal (Figure 3, 3^rd^ row). These results confirm that most of the nascent flagellar proteins are incorporated into the axoneme from the distal tip and that the proteins undergo little turnover once assembled.

**Figure 3.**
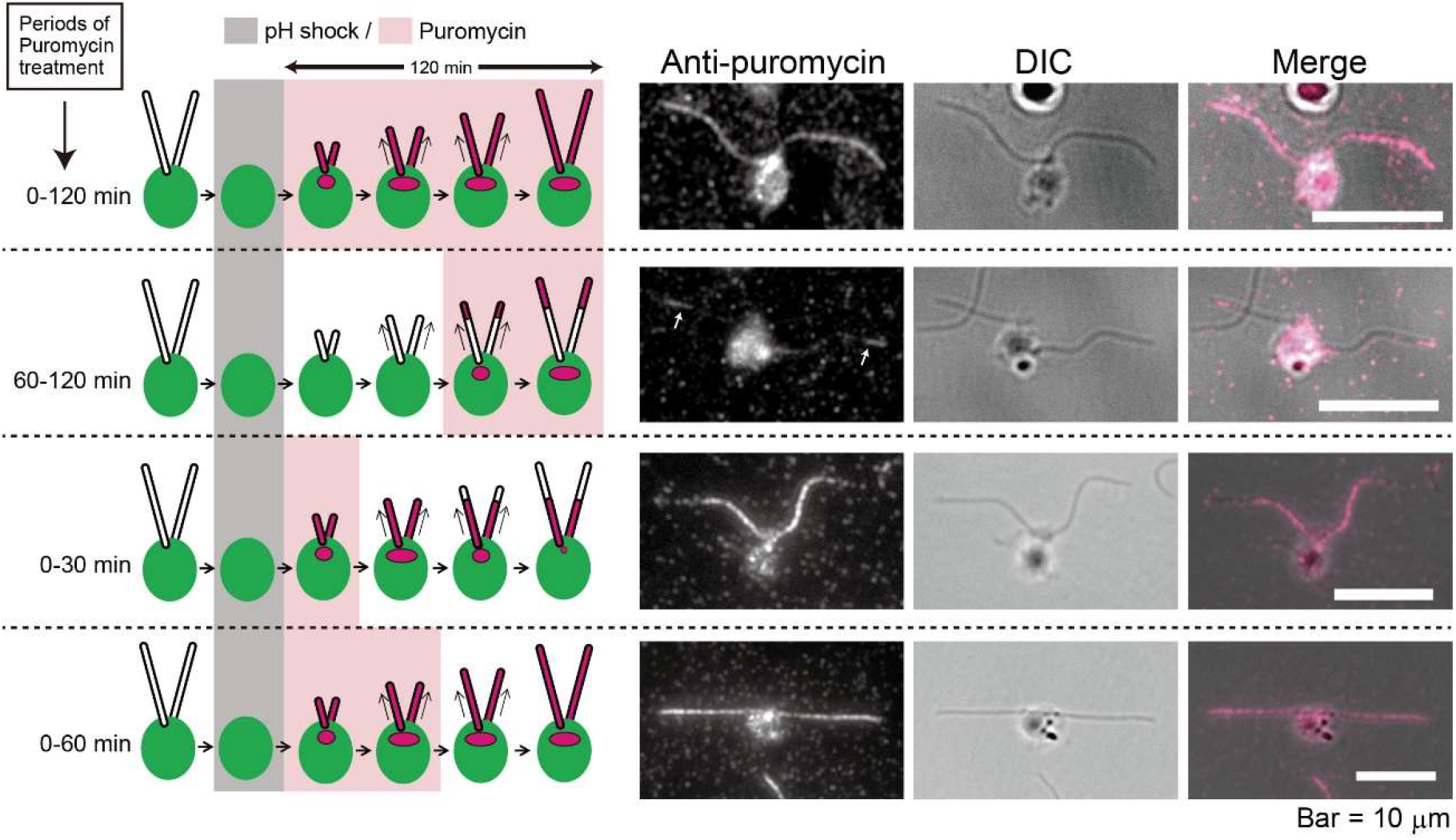
Newly synthesized proteins are incorporated from the tip of the flagellum. Indirect immunofluorescence microscopy of cells after flagellar regeneration. Schematic illustrations on the left panels indicate the timing of puromycin application during flagellar regeneration.

### Puromycin incorporation into flagella is tightly coupled with protein synthesis

We next examined the effect of another protein synthesis inhibitor, cycloheximide, to determine whether the observed flagellar incorporation of puromycin is in fact coupled with the synthesis of flagellar proteins. We found 1 μg/ml cycloheximide prevented cellular uptake of puromycin (Figure 4A) as well as flagellar assembly beyond half the normal length, consistently with previous studies (Rosenbaum et al, 1969; Lefebvre et al, 1978; Figure 4B). This observation strongly suggests that the flagellar incorporation of puromycin requires protein synthesis.

**Figure 4.**
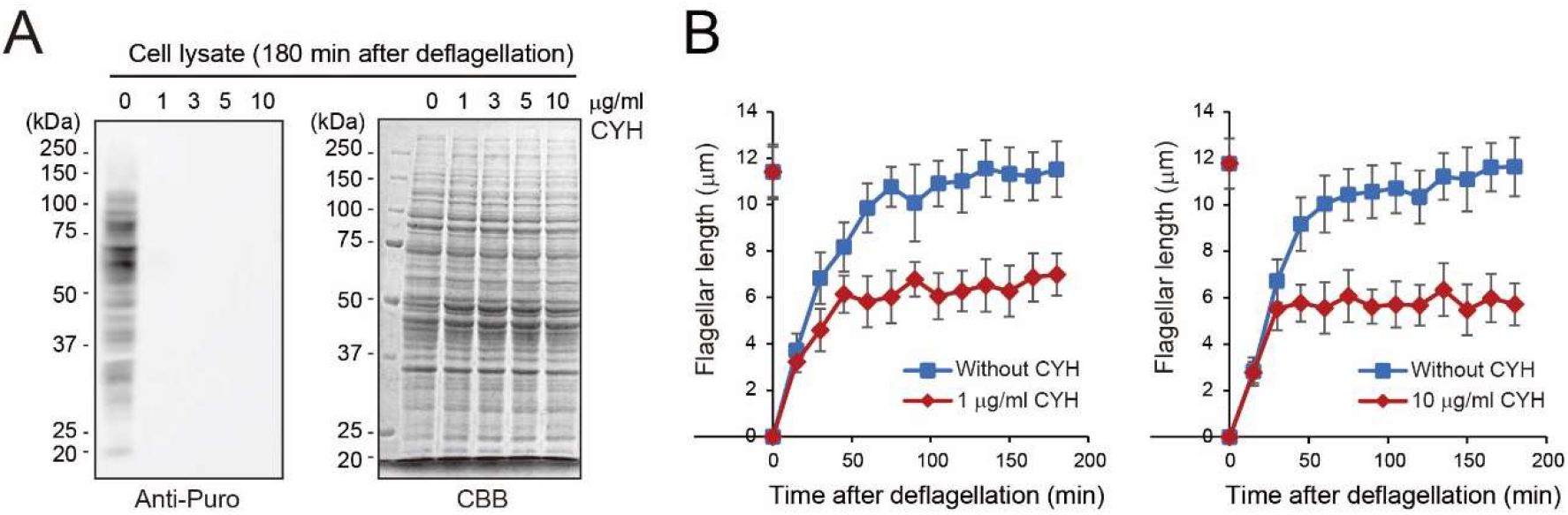
Cycloheximide almost completely inhibits protein synthesis as determined by SUnSET. (A) Western blotting and CBB-stained gel of wild-type cells after flagellar assembly. Deflagellated cells were treated with cycloheximide (0, 1, 3, 5, or 10 μg/ml) in the presence of 50 μg/ml puromycin for 180 min. (B) Flagellar regeneration kinetics of the cells with or without cycloheximide.

To rule out a small but nonnegligible possibility that cycloheximide prevented puromycin incorporation through a protein synthesis-independent mechanism, we next used a cycloheximide-resistant mutant, *act2*, which carries a proline to leucine substitution in the ribosomal protein RPL36a (Fleming et al, 1987; Stevens et al, 2001; Supplemental Figure 1A). While this mutant was almost indistinguishable from wild type in growth rate and flagellar regeneration kinetics in normal media (Supplemental Figure 1B, 2A), the two strains strikingly differed in cycloheximide sensitivity; in contrast to wild type, *act2* regenerated almost full-length flagella in the presence of 10 μg/ml cycloheximide (Supplemental Figure 2A). When 50 μg/ml puromycin was applied to the cells undergoing flagellar regeneration in the cycloheximide-containing medium, puromycin uptake occurred normally in *act2* but not in wild type (Supplemental Figure 2B). Consistent with these results, immunofluorescence observation of NFAps showed that *act2* cells treated with cycloheximide and puromycin displayed normal-length flagella with puromycin signals along the length, while the wild type cells treated in the same manner yielded short flagella essentially devoid of signals (Supplemental Figure 2C). These results establish that the *act2* mutant can synthesize proteins in the presence of 10 μg/ml cycloheximide, and that the flagellar incorporation of puromycin indeed represents newly synthesized proteins.

### Why were apparently complete proteins labeled with puromycin?

The observation that puromycin-labeled proteins tended to localize along the flagellar length in a dotted manner (Figure 2A) may indicate that only a small portion of newly synthesized proteins were labeled by puromycin. The exact puromycin-labeled fraction of all cellular and flagellar proteins at a given puromycin concentration remains to be quantified. Western blotting of flagellar proteins from puromycin-treated cells displayed numerous bands (Figure 2B). Most notably, these bands appeared almost identical in mobility to the bands in the silver-stained gel, suggesting that incompletely translated (C-terminally truncated) products were little incorporated into the flagellar sample. The apparent absence of incomplete proteins in flagella may be due to a rapid degradation of incomplete proteins in the cytoplasm, inability of incomplete proteins to assemble into the axoneme, or preferential addition of puromycin to the C-termini of complete products. The true reason for the absence of incomplete axonemal proteins also needs to be explored by examining puromycin-labeled cytoplasmic proteins.

Although several questions have remained to be answered in the future studies, our present study clearly showed that the SUnSET method can be applied to examine flagellar assembly as a useful tool to visualize newly synthesized proteins. Preferential puromycin incorporation into full-length proteins appears to make this method particularly valuable.

## Materials and Methods

### Strains and cultures

The C. *reinhardtii* strains used in this study were wild type (cc124, cc125) and *act2* (CC-1590, mt-; obtained from the *Chlamydomonas* resource center). The *act2* mutant was backcrossed twice to wild-type strains before the analyses. Cells were routinely grown and maintained on Tris-acetate-phosphate (TAP; Gorman and Levine, 1965) agar plates. For experiments, aliquots of cells on the plate were transferred to TAP liquid medium with aeration. Cells were grown on a 12-12 h light-dark cycle with 25°C. Optical densities (750 nm) of cell cultures were measured each day to track the cell growth.

### Flagellar regeneration

To deflagellate cells, the pH of liquid cultures was decreased and maintained at 4.5 for 20 s by the dropwise addition of 0.5 M CH_3_COOH. Subsequently, the pH was returned to 7.4 by the dropwise addition of 0.5 M KOH. After deflagellation, aliquots of cells were isolated and fixed with 1% glutaraldehyde at 15 min intervals up to 180 min. Flagella from at least 30 cells were measured by ImageJ to obtain their average lengths.

### Isolation and fractionation of flagella

Flagella were isolated according to Witman et al (1972) with some modification. Isolated flagella were suspended in HMDEK buffer (30 mM Hepes, 5 mM MgSO_4_, 1 mM DTT, 1 mM EGTA, 50 mM CH_3_COOK). For fractionation, flagella were frozen and thawed twice with liquid nitrogen to obtain a supernatant (matrix fraction; 25,000 g, 10 min, 4°C) (Fan et al, 2010). Pelleted flagella were subsequently resuspended with HMDEK containing 0.1% Igepal CA-630 (Sigma Aldrich; an alternative for Nonidet-40) and placed on ice for 10 min to release membrane. After a centrifugation (25,000 g, 10 min, 4°C), the supernatant (membrane fraction) and the pellet (axonemal fraction) were isolated.

### Surface sensing of translation (SUnSET) method

Unless stated otherwise, cells in the TAP liquid culture were treated with 50 μg/ml puromycin (P8833; Sigma Aldrich) for indicated periods. Flagella from puromycin-treated cells were isolated as described above. For the whole cell sample, cells were collected with gentle centrifugation (2,000 g, 3 min) and washed twice with 10 mM Hepes. The cells were then suspended in HMDEK, frozen and thawed twice with liquid nitrogen. Whole cell sample was obtained as a supernatant after a centrifugation (25,000 g, 10 min, 4°C). Puromycin-labeled proteins were detected either by Western blotting or by indirect fluorescence microscopy using a monoclonal anti-puromycin antibody (12D10; Sigma Aldrich).

### SDS-PAGE and Western blotting

Proteins were separated by 9% SDS-polyacrylamide gel electrophoresis and blotted on a PVDF membrane (Towbin et al, 1979). Anti-puromycin (12D10; Sigma) and anti-α-tubulin (B512; Sigma) antibodies were used.

### Indirect immunofluorescence microscopy

Indirect immunofluorescence microscopy was performed according to Kubo et al (2015). Cells were treated with autolysin (Picariello et al, 2020) to remove cell walls and attached to a microscope slide treated with 0.01% polyethylenimine. Nucleoflagellar apparatuses (NFAps) were obtained by vigorously treating the cells with 1% Igepal CA-630 (Sigma Aldrich) (Sanders and Salisbury, 1995). The NFAps were fixed with 1% paraformaldehyde and were treated with cold methanol and acetone (−20 °C) for 5 min respectively. The NFAp were incubated with primary antibodies for 1 h followed by secondary antibodies conjugated with either Alexa488 or Alexa594 (Invitrogen). SlowFade Gold Antifade Mountant (Invitrogen) was used to reduce fading of fluorescence. The samples were observed by a fluorescence microscope (BX53; Olympus) and images were collected using a CCD camera (ORCA-Flash4.0; Hamamatsu PhotonICs).

## Acknowledgement

We greatly appreciate Dr. Ritsu Kamiya (Chuo University, Tokyo) for critically reading the manuscript and a constructive discussion. We also appreciate Dr. Keigo Morita (The University of Tokyo) and Dr. Yoshitaka Kurikawa (The University of Tokyo) for technical advice. This work was supported by Takeda Science Foundation (to T.K. and T.O.), the Uehara Memorial Foundation (to T.K.) and Japan Society for the Promotion of Science [17K15115 (to T. K.), 19K16123 (to T.K.), and 21H02654 (to T.O.)].

## Author contributions

T.K. designed the research; T.K. and N.K. performed the experiments; T.K. and T.O. analyzed data; and T.K. and T.O. wrote the paper. All authors reviewed the manuscript.

## Corresponding authors

Correspondence to Tomohiro Kubo and Toshiyuki Oda.

---

**Supplemental figure 1.**
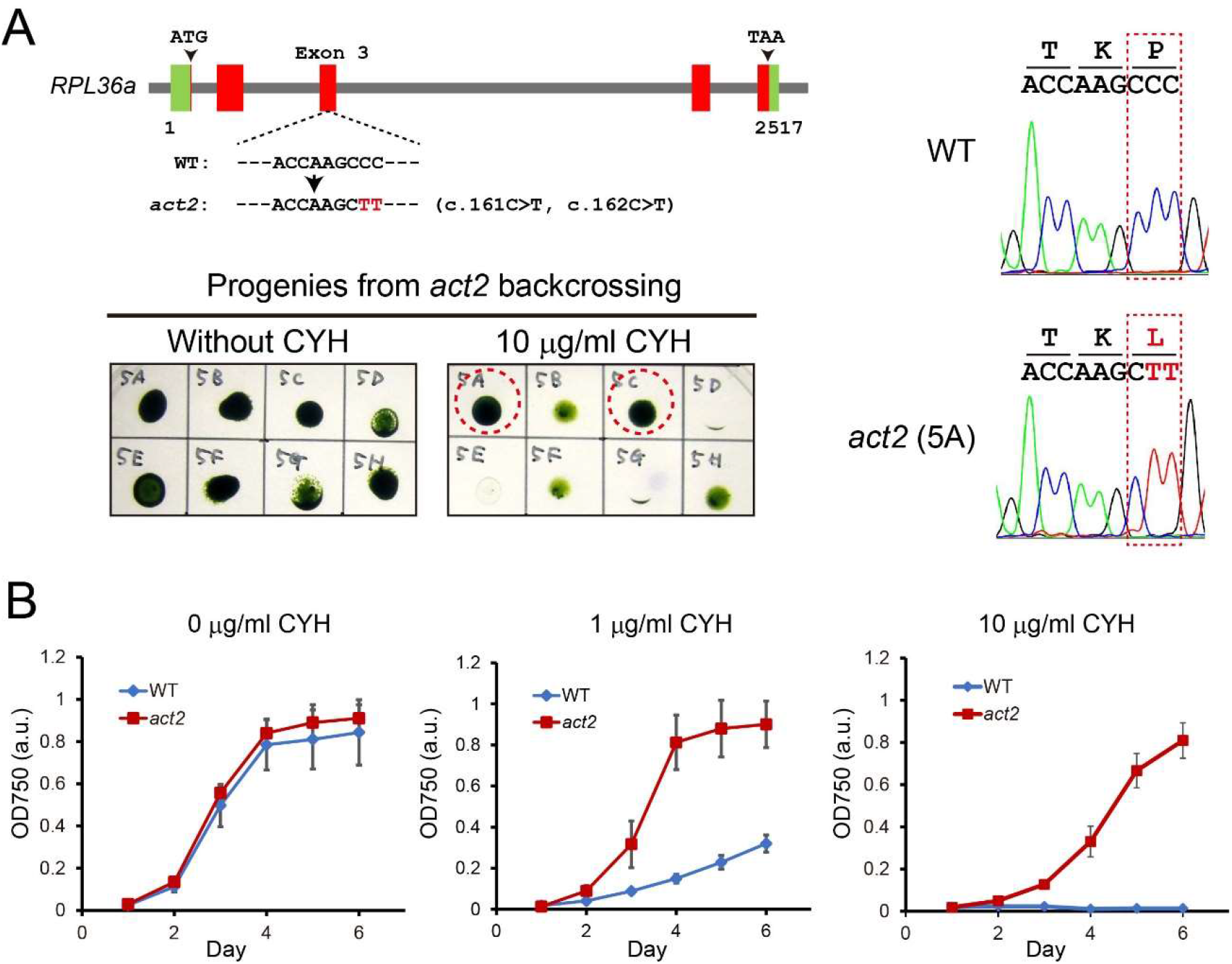
Characterization of the mutant *act2* resistant to cycloheximide. (A) Structure of *RPL36a* gene (Cre06.g310700) (upper). The mutant *act2* possesses proline to leucine substitution due to “CCC” to “CTT” mutation. The original *act2* (CC-1590) was backcrossed twice with wild type. The obtained progenies were cultured with or without 10 μg/ml cycloheximide (below). Genomic sequencing of clone 5A (right). (B) Proliferation rates of wild-type and *act2* cells in the presence of 0, 1, or 10 μg/ml cycloheximide. The daily cell densities (OD750) were measured.

**Supplemental figure 2.**
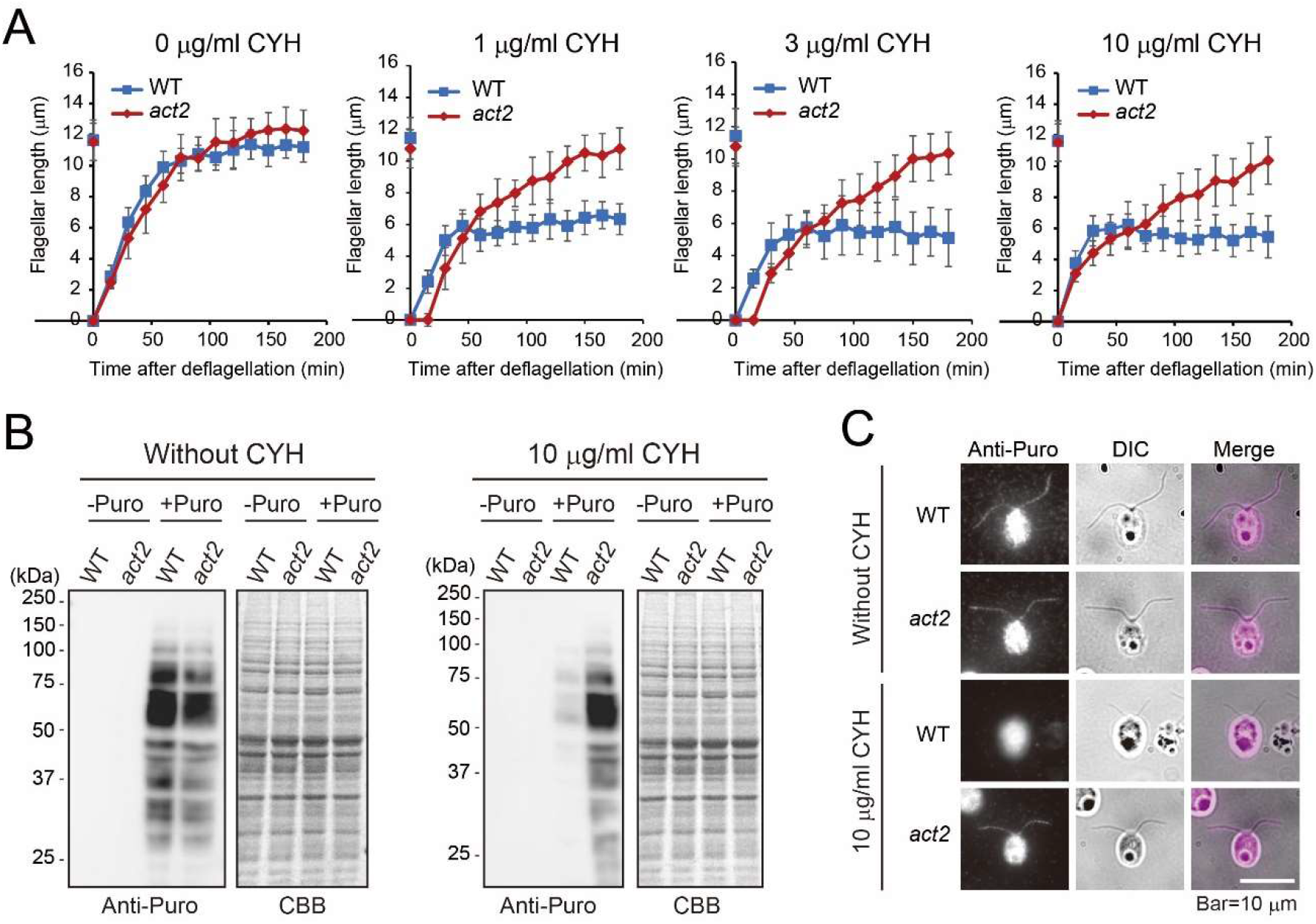
The mutant *act2* regenerates full-length flagella after pH shock-induced deflagellation in the presence of cycloheximide. (A) Flagellar regeneration of wild type and *act2* in the presence of cycloheximide (0, 1, 3, or 10 μg/ml). (B) Puromycin uptake of wild-type and *act2* cells with (right) or without (left) 10 μg/ml cycloheximide. Deflagellated cells were treated with or without 50 μg/ml puromycin for 180 min. (C) Immunofluorescence observation of wild type and *act2* with regenerated flagella. The cells were treated with or without 10 μg/ml cycloheximide in the presence of 50 μg/ml puromycin.

## References

Corbit KC, Aanstad P, Singla V, Norman AR, Stainier DY, and Reiter JF. (2005). Vertebrate Smoothened functions at the primary cilium. Nature. 437, 1018–1021.

Fan ZC, Behal RH, Geimer S, Wang Z, Williamson SM, Zhang H, Cole DG, and Qin H. (2010). *Chlamydomonas* IFT70/CrDYF-1 is a core component of IFT particle complex B and is required for flagellar assembly. Mol Biol Cell. 21, 2696–2706.

Fleming GH, Boynton JE, and Gillham NW. (1987). The cytoplasmic ribosomes of *Chlamydomonas reinhardtii:* characterization of antibiotic sensitivity and cycloheximide-resistant mutants. Mol Gen Genet. 210, 419–428.

Gorman DS and Levine RP. (1965). Cytochrome f and plastocyanin: their sequence in the photosynthetic electron transport chain of *Chlamydomonas reinhardti*. Proc. Natl. Acad. Sci. USA 54, 1665–1669.

Huangfu D, Liu A, Rakeman AS, Murcia NS, Niswander L, and Anderson KV. (2003). Hedgehog signaling in the mouse requires intraflagellar transport proteins. Nature. 426, 83–87.

Johnson KA, and Rosenbaum JL. (1992). Polarity of flagellar assembly in *Chlamydomonas*. J Cell Biol. 119, 1605–1611.

Kozminski KG, Johnson KA, Forscher P, and Rosenbaum JL. (1993). A Motility in the Eukaryotic Flagellum Unrelated to Flagellar Beating. Proc Natl Acad Sci U S A. 90, 5519–5523.

Kubo T, Hirono M, Aikawa T, Kamiya R, and Witman GB. (2015). Reduced tubulin polyglutamylation suppresses flagellar shortness in *Chlamydomonas*. Mol Biol Cell. 26, 2810–2822.

Lefebvre PA, Nordstrom SA, Moulder JE, and Rosenbaum JL. (1978). Flagellar elongation and shortening in *Chlamydomonas*. IV. Effects of flagellar detachment, regeneration, and resorption on the induction of flagellar protein synthesis. J Cell Biol. 7, 8–27.

Marshall WF, and Rosenbaum JL. (2001). Intraflagellar transport balances continuous turnover of outer doublet microtubules: implications for flagellar length control. J Cell Biol. 155, 405–414.

Nathans D. (1964). Puromycin inhibition of protein synthesis: incorporation of puromycin into peptide chains. Proc Natl Acad Sci U S A. 51, 585–592.

Picariello T, Hou Y, Kubo T, McNeill NA, Yanagisawa HA, Oda T, and Witman GB. (2020). TIM, a targeted insertional mutagenesis method utilizing CRISPR/Cas9 in *Chlamydomonas reinhardtii*. PLoS One. 15, e0232594.

Piperno G, Mead K, Henderson S. (1996). Inner arm but not outer dynein arms require the activity of kinesin homologue protein KHP1 (FLA10) to reach the distal part of flagella in *Chlamydomonas*. J Cell Biol. 133, 371–379.

Rosenbaum JL, and Child FM. (1967). Flagellar regeneration in protozoan flagellates. J Cell Biol. 34, 345–364.

Rosenbaum JL, Moulder JE, Ringo DL. (1969). Flagellar elongation and shortening in Chlamydomonas. The use of cycloheximide and colchicine to study the synthesis and assembly of flagellar proteins. J Cell Biol. 41, 600–619.

Rosenbaum JL, and Witman GB. (2002). Intraflagellar transport. Nat Rev Mol Cell Biol. 3, 813–825. Review.

Sanders MA, and Salisbury JL. (1995). Immunofluorescence microscopy of cilia and flagella. Methods Cell Biol. 47, 163–169.

Schmidt E.K, Clavarino G, Ceppi M, and Pierre P. (2009). SUnSET, a nonradioactive method to monitor protein synthesis. Nat Methods. 6, 275–277.

Song L, and Dentler WL. (2001). Flagellar protein dynamics in *Chlamydomonas*. J Biol Chem. 276, 29754–29763.

Stevens DR, Atteia A, Franzén LG, and Purton S. (2001). Cycloheximide resistance conferred by novel mutations in ribosomal protein L41 of *Chlamydomonas reinhardtii*. Mol Gen Genet. 264, 790–795.

Towbin H, Staehelin T, and Gordon J. (1979). Electrophoretic transfer of proteins from polyacrylamide gels to nitrocellulose sheets: procedure and some applications. Proc. Natl. Acad. Sci. USA 76, 4350–4354.

Witman GB, Carlson K, Berliner J, and Rosenbaum JL. (1972). *Chlamydomonas* flagella. I. Isolation and electrophoretic analysis of microtubules, matrix, membranes, and mastigonemes. J Cell Biol. 54, 507–539.

